# Convex Optimized Population Receptive Field (CO-pRF) Mapping

**DOI:** 10.1101/172189

**Authors:** David Slater, Lester Melie-Garcia, Stanislaw Adaszewski, Kendrick Kay, Antoine Lutti, Bogdan Draganski, Ferath Kherif

## Abstract

Population receptive field (pRF) mapping represents an invaluable non-invasive tool for the study of sensory organization and plasticity within the human brain. Despite the very appealing result that fMRI derived pRF measures agree well with measurements made from other fields of neuroscience, current techniques often require very computationally expensive non-linear optimization procedures to fit the models to the data which are also vulnerable to bias due local minima issues. In this work we present a general framework for pRF model estimation termed *Convex Optimized Population Receptive Field* (*CO*-*pRF*) mapping and show how the pRF fitting problem can be linearized in order to be solved by extremely fast and efficient algorithms. The framework is general and can be readily applied to a variety of pRF models and measurement schemes. We provide an example of the CO-pRF methodology as applied to a computational neuroimaging approach used to map sensory processes in human visual cortex - the CSS-pRF model. Via simulation and in-vivo fMRI results we demonstrate that the CO-pRF approach achieves robust model fitting even in the presence of noise or reduced data, providing parameter estimates closer to the global optimum across 93% of in-vivo responses as compared to a typical nonlinear optimization procedure. Furthermore the example CO-pRF application substantially reduced model fitting times by a factor of 50. We hope that the availability of such highly accelerated and reliable pRF estimation algorithms will facilitate the spread of pRF techniques to larger imaging cohorts and the future study of neurological disorders and plasticity within the human brain.

**Highlights:**

- Adaptable to an arbitrary computational pRF model, sensory modality and imaging modality.
- Model fitting accelerated by up to a factor of 50.
- CO-pRF parameter estimates achieve a better fit, closer to the global optimum, as compared to typical nonlinear implementations which suffer from local minima issues.
- Robust parameter estimation even at low CNR and reduced scan time.

## Introduction

A principle goal in sensory neuroscience is to accurately predict neural responses to a wide range of sensory stimuli. For example, one would like to predict the way in which any visual input generates neural activity across the visual system. For the auditory system one would like to predict the way in which auditory stimuli of a given spectral profile generate neural responses in auditory cortex. Though there remains much work in order to achieve this goal across human sensory cortex, a number of advances have been made to produce models that directly predict how the human brain responds to sensory stimuli.

The last two decades has seen a number of non-invasive fMRI techniques being developed for human visual field mapping (DeYoe et al., 1996; Engel et al., 1994; Sereno et al., 1995) and more recently for population receptive field (pRF) mapping (Dumoulin and Wandell, 2008). Both of these methods provide estimates of the visual field position that generates the maximal response in a voxel by modeling the fMRI time-course for a set of known visual stimuli. In addition the pRF technique provides quantitative estimates of the visual field area over which a voxel responds, its *pRF size.* Within the auditory system recent work has begun to investigate the full frequency response profile of auditory cortex voxels (De Martino et al., 2013; Moerel et al., 2012). In analogy to the visual system pRF methodology of Dumoulin and Wandell (2008) recent work by Thomas et al. (2015) developed a computational pRF model relating the spectral profiles of auditory stimuli to auditory receptive field properties and their generated BOLD fMRI time series. The method is able to estimate the preferred response frequency of an auditory voxel as well as the tuning width of frequencies over which it responds.

The visual and auditory system pRF methodologies have the advantage that they incorporate the use of general models that have the ability to predict responses for a wide range of stimulus sequences. Furthermore pRF techniques provide quantitative estimates of neural population receptive field properties which can be compared across different measurement instruments and species. The key difference between pRF modeling and earlier methods is that pRF techniques apply an explicit computational model of the fMRI response in terms of input-referred parameters (Wandell and Winawer, 2015).

The strength and flexibility of the pRF technique has resulted in a number of extensions to the original visual pRF model. Zuiderbaan et al. (2012) suggested modeling the receptive field with two rather than one Gaussian. The second Gaussian models the negative surround effect and the systematic drop bellow baseline signal levels. A number of other authors have suggested modeling receptive fields using different shapes (Greene et al., 2014; Kay et al., 2008; Lee et al., 2013; Sprague and Serences, 2013). Another extension to the pRF model is the compressive spatial summation (CSS) model (Kay et al., 2013a) which includes a compressive static nonlinearity applied after stimulus summation. This extension considerably improved the number of visual stimuli the model could accurately predict and revealed differences in spatial summation across visual field maps. For a detailed review of visual system pRF applications see Wandell and Winawer (2015).

Despite the power of pRF methods to map sensory brain systems and estimate quantitative response properties the non-linear optimization routines typically used to fit such models are extremely computationally intensive. This causes practical problems when applying these techniques to clinical cohorts or longitudinal studies with large scan numbers. Additionally, non-linear fitting algorithms are highly sensitive to the initial seed parameters and often stop at local-rather than global-minima. This introduces a potential bias as parameter estimates may be dependent upon the choice of initial seed location, introducing inaccuracy and unwanted variance to pRF parameter estimates.

Here we present a new framework for estimating pRF measures by taking advantage of convex optimization techniques which we call *CO*-*pRF* standing for *Convex Optimized Population Receptive Field* mapping. The pRF model is reformulated into an equivalent linearized problem which can be solved using highly efficient algorithms. The CO-pRF framework is general and permits the potential estimation of pRF parameters across any sensory modality, imaging technique or computational model form. We first present the generalized CO-pRF framework and then as a proof of principle we demonstrate the potential gains in model estimation time, precision and accuracy when CO-pRF is applied to the CSS-pRF model of visual function (Dumoulin and Wandell, 2008; Kay et al., 2013). However we note that the flexibility of the CO-pRF estimation approach makes it trivial to linearize additional computational pRF models of an arbitrary form.

The CO-pRF methodology and design is described in the following sections. We present the results of numerical simulations and in-vivo fMRI data acquisitions, discussing the benefits, flexibility and potential hazards of the CO-pRF framework. A Matlab toolbox for applying CO-pRF to a variety of pRF models is made available at https://github.com/davesl/COpRF. We hope that this toolbox will be useful to the sensory neuroscience community and plan to expand upon its development over the coming years.

## Materials and Methods

Here we first present the pRF model (Dumoulin and Wandell, 2008) and show how this can be reformulated into a linearized system of equations which can be solved efficiently using convex optimization techniques. This is the generalized CO-pRF framework which can be readily adapted to any pRF model. We then discuss the application of the framework to the specific case of the extended CSS-pRF model (Kay et al., 2013).

### A generalized CO-pRF framework

For a pRF experiment of arbitrary design the measured physiological signal is related to the receptive fields as follows:

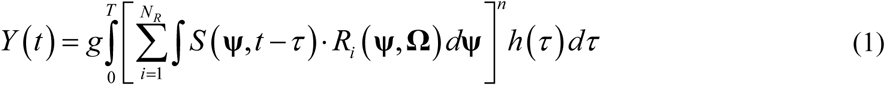

where *Y* (*t*) represents the physiological measurement signal, *S* (**ψ**, *t* − *τ*) is the stimulus function, **ψ** is the set of parameters **ψ** = [*ψ*_1_,…, *ψ_N_*] defining the stimuli of the experiment. In the case of a visual pRF experiment this would be the position of the pixel **ψ** = [**r⃗**, *A*] = [*x*, *y*, *A*] in stimulus space that is on at time *t* and its luminance, *A*, (or amplitude), though **ψ** could also include additional stimulus parameters such as the distribution of second order contrast (Kay et al., 2013b) or chromatic information of the visual stimulus. For the case of auditory pRF mapping **ψ** could be the set of frequencies described by spectral profiles of auditory stimuli presented at time *t*, **ψ** = [*ω*_1_,…,*ω_N_*]. *R_i_* (**ψ**, **Ω**) defines the pRF function for the *i*-th measured response which evidently depends on the stimulus parameters **ψ** as well as the intrinsic parameters **Ω** defining the form of this function at a particular measurement locus. For instance if applied to the original visual pRF study of Dumoulin and Wandell (2008) **Ω** would describe a circularly symmetric Gaussian function that is defined by a receptive field center and width. Though in practice there is no reason why **Ω** might not be used to model more complicated receptive field shapes (e.g. Zuiderbaan et al., 2012). Typically one would assume that *R_i_* is time invariant over the time course of a typical experiment. However in the future it could be possible to introduce parameters which model the plasticity of receptive fields over time or the influence of attention on receptive field properties (Kay et al., 2015; Sprague and Serences, 2013). The CO-pRF framework is therefore highly flexible such that *S* and *R_i_* may take any form. Though it is noted that *S* should be carefully constructed to reduce degeneracy in the response forms of receptive fields with different underlying parameters (e.g. for visual pRF mapping, stimuli should be capable of probing both polar angle and eccentricity).

In the generalized expression of Eq. 1 it is assumed that several receptive fields might coexist within a measurement locus. This would be expected for fMRI experiments where the spatial resolution is in the order of millimeters. This is indicated by the sum across the *N_g_* number of possible families of receptive field functions.

The neuronal signal coupling function is described using *h* (*τ*) which models across a total duration of time *T*. For the case of fMRI signal coupling *h* (*τ*) takes the form of the well-known hemodynamic response function (HRF). If working with local field potential (LFP) data *h* (*τ*) could represent the neural mass model of signal coupling (David and Friston, 2003). The exponent parameter *n* can be used to perform a compressive summation across the stimuli (Kay et al., 2013a) or left equal to 1 for the more typical case of linear summation. Finally *g* describes the scaling factor, or gain, of the measurement system.

Typically non-linear fitting algorithms are used to estimate the pRF parameters by minimizing residual error. However, as the response functions can be estimated a priori using Eq. (1) we can reformulate the measurement process as the following system of linear equations:

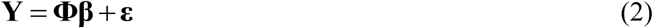

where **Y** ∈ ℝ^*N_t_*^ is the vector containing the *N_t_* physiological measurement signals, **Φ** ∈ ℝ^*N_t_*×*N_s_*^ is the linear operator (or dictionary) of *N_s_* modeled response functions (also called atoms),
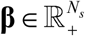
are the dictionary
coefficients to be estimated and **ε** models the acquisition noise not accounted for by the model. Thus the linear problem of Eq. (2) can be solved using a variety of algorithms based on convex optimization. Without a loss of generality the existing methods can be expressed with the following general regularized least-squares formulation:

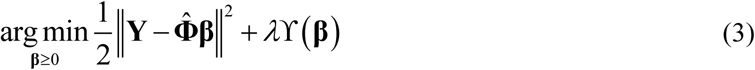

where **∥ ∥** is the standard *l*_2_-norm and the positivity constraint **β** ≥ 0 is explicitly imposed as the coefficients **β** correspond to the modeled response functions contained in the dictionary **Φ**. **Φ̂** denotes the dictionary after each column has been normalized by the standard *l*_2_-norm. The operator ϒ(·) represents a generic regularization function and *λ* > 0 controls the trade-off between data fitting and regularization terms. An optimal value for *λ* can either be set empirically or via a data driven approach (Golub et al., 1979; Hansen, 2000). Eq. (3) is reduced to the standard non-negative least-squares regression when *λ* = 0. However in most cases a regularization term is required in order to promote stability of the solution or to introduce prior knowledge. A common regularization approach for promoting sparsity in the estimated dictionary coefficients is via the standard *l*_1_-norm, **Ψ** = **∥ ∥**_1_. This has been used when solving various inverse problems related to tissue microstructure in diffusion MRI (Daducci et al., 2014; Landman et al., 2012; Michailovich et al., 2010; Ramirez-Manzanares et al., 2007).

### CO-pRF applied to the CSS-pRF model

Here we describe an example application of the CO-pRF framework to the modeling of fMRI responses in human visual cortex. We chose the CSS-pRF model of Kay et al. (2013) as it has been shown to generalize well to a broader range of stimuli than simpler models which assume linear spatial summation and was recently used to assess the effect of attention on pRF properties in ventral temporal cortex (Kay et al., 2015). Our choice of a visual system pRF model reflects the fact that models of this type are better established within the literature and more widely applied than pRF techniques in other sensory modalities (e.g. auditory pRF models; Thomas et al., 2015).

The CSS-pRF method models fMRI time series responses by computing the integral over stimulus space of a circularly symmetric Gaussian receptive field intersecting with the stimuli at time *t* and then applying a static power-law nonlinearity (Kay et al., 2013a). This is expressed via the following adaptation of Eq. (1):

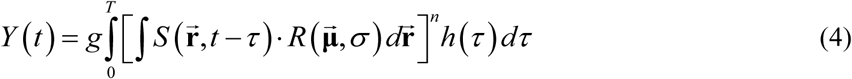

where **ψ** = [**r⃗**, *A*)] = [*x*, *y*, *A*] defines the Euclidian position of all pixels in stimulus space along *x* and *y* axes with limits [*x*_0_,*x_S_*] and [*y*_0_,*y_S_*] respectively. The amplitude *A* is normally assumed to be binarized e.g. *A*=1 when a stimulus is being displayed through an aperture window. We assume the same function for all receptive fields within a voxel defined by the function *R* and described by a circularly symmetric Gaussian with mean **μ⃗** = [*μ_x_*, *μ_y_*] and standard deviation *σ*. Our computational model of fMRI responses is thus described by the following equation:

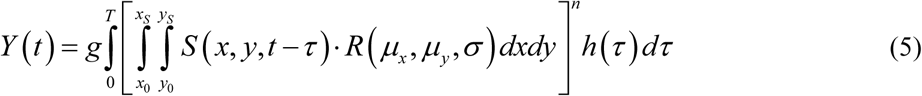

When reformulating the CSS-pRF model of Eq. (5) into the CO-pRF framework we first design a receptive field dictionary, **D** ∈ ℝ^*N_v_*×*N_s_*^, with *N_s_* combinations of the model parameters by *N_v_* pixels where the receptive fields are spatially represented. In the case of the CSS-pRF model we have *p*=5 unknown model parameters to estimate: x-position (*μ_x_*), y-position (*μ_y_*), pRF size (*σ*), compressive exponent (*n*) and gain (*g*). The Cartesian coordinate system of *x* and *y* can be expressed equivalently in polar coordinates using polar angle (*θ*) and eccentricity (*r*) parameters.

The final response dictionary, **Φ** ∈ ℝ^*N_t_*×*N_s_*^, is then estimated as a combination of **D** and the specific stimulus pattern **S** ∈ ℝ^*N_t_*×*N_v_*^ with *N_t_* time points and *N_v_* pixels as [**S · D**]^*n*^ convoluted by the hemodynamic response along the time dimension and multiplied by *g*. The *i*-th column in **Φ** corresponds to the predicted fMRI time series modeled using the *i*-th set of pRF parameters used to construct **D** interacting with the stimuli S. After constructing **Φ** we can reformulate the CSS-pRF model as a convex optimization problem:

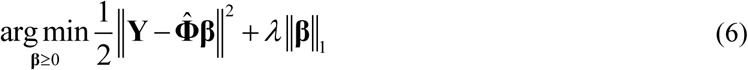

where ϒ = **∥ ∥**_1_ is the standard *l*_1_-norm and employed to promote sparsity in the dictionary coefficients estimated via the regression.

In this work we adopted the following implementation to build *Φ* and estimate the CSS-pRF model parameters in an efficient manner.

- The computational costs required to estimate the **β** coefficients are dependent upon the total number of atoms in the dictionary, *N_s_*. To improve estimation efficiency we implemented a hierarchical fitting procedure. We first estimate the pRF parameters on a reduced size sub-dictionary, **Φ**_1_, to localize the approximate *x* and *y* pRF position. We then select atoms from a dense response dictionary, **Φ**_2_, which have *x* and *y* position parameters within a given radius of the initial sub-dictionary estimates. In this work the radius was set to be 10% of the total stimulus coverage.
- The gain parameter, *g*, acts as a simple linear scaling factor on the modeled responses. We do not need to model this explicitly and can instead calculate the gain from **β** and the normalization factors used to normalize the dictionary (see Eq.(7) and (8)). This reduces the final dictionary size considerably and the number of parameters we need to explicitly model and estimate to 4.
- When constructing the dense dictionary **Φ**_2_ we chose a set of underlying parameters for **D** ∈ ℝ^*N_p_*×*N_s_*^ which covered the full stimulus area and expected range of pRF parameters. Specifically for the *x* and *y* pRF positions we create a regular Cartesian grid extending out to 120% of the angular stimulus coverage for a total of *N_xy_* = 1804 *x* and *y* parameter combinations. We considered 10 values for *σ* ∈ {l,…,64} and 6 values for *n* ∈ {0.1,…,1}. Thus we have a total dictionary size of *N_s_* = *N_xy_* × *N_σ_* × *N_n_* = 1804 × 10 × 6 = 108240 atoms.
- The reduced size sub-dictionary **Φ**_1_ is constructed in the same manner as for **Φ**_2_, over the same range but with a reduced number of parameters. The total number of parameters used in **Φ**_1_ was *N_s_* = *N_xy_* × *N_σ_* × *N_n_* = 293 × 4 × 3 = 3516 atoms.
- For both dictionaries **Φ**_1_ and **Φ**_2_ we remove any atoms for which *σ* does not overlap with the stimulus area. This removes atoms that have little or no response to the given stimuli.
- It has been shown that enforcing a sparsity prior as a regularization term can result in biased solutions because the *l*_1_-norm will often underestimate the genuine **β** coefficients (Figueiredo et al., 2007). Thus every time we solve Eq. (6) we apply a debiasing step by solving Eq. (6) once more, this time using only the atoms with non-zero **β** coefficients and without regularization (e.g. *λ* = 0). This additional step acts to correct the estimated weights in *β* (Daducci et al., 2014).
- After solving Eq. (6) and the following debiasing step we can estimate the model parameters from the **β** coefficients via the following model inversion:

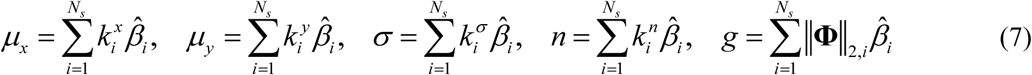

where,

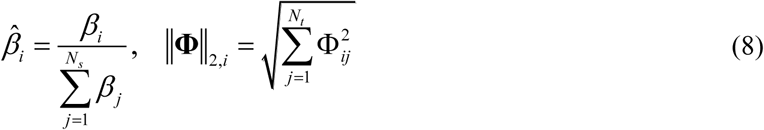

for all *i* ∈ {1, …, *N_s_*} we denote
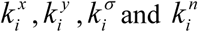
respectively as the x-position, y-position, pRF size, and the compressive exponent corresponding to the *i*-th atom in **Φ**. ∥**Φ**∥_2,*i*_ = ℝ^*N_s_*^ is the *l*_2_-norm of the *i*-th atom which was used to form the normalized response dictionary, **Φ̂**, before solving Eq. (6). The procedure takes the weighted sum of underlying pRF model parameters to convert a set of multiple atoms into a single model solution. This assumes that the response function is continuous over the range of parameters modeled within the dictionary. Such methods have been used to accelerate and improve parameter estimation in diffusion MRI models of tissue microstructure (Daducci et al., 2014).
- We solve Eq. (6) over a range of ever decreasing values for *λ*. For each iteration we use the estimated pRF parameters of Eq. (7) and (8) and the generative CSS-pRF model of Eq. (5) to calculate the prediction error between modeled and real data time series. Thus model fitness is determined directly from the parameter estimates and not the weighted-sum of atoms in Eq. (2). We stop when we observe an increase in model prediction error between successive iterations. This procedure gradually relaxes the sparsity constraint, allowing ever more atoms to contribute to **β** until the model parameter explanatory power is reduced. This approach was chosen to optimize efficiency, however alternatives such as generalized cross-validation (Golub et al., 1979) or L-curve optimization (Hansen, 2000) techniques could equally be used to select *λ*. We will refer to this convex reformulation of the CSS-pRF model as CSS_CO-pRF_ as opposed to the original version which we shall call CSS_Orig_.

### Stimulus description

Visual stimuli were created using extensions from the PsychToolbox (Brainard, 1997; Pelli, 1997) within the Matlab programming environment. An LCD projector was used to image the stimuli onto a back projection screen within the bore of the magnet. Subjects viewed the display through an angled mirror. The maximal stimulus radius was 9.15° of visual angle. Each stimulus image was at a resolution of 768 × 768 pixels. For all experiments a small dot at the centre of the image served as a fixation point (4 pixel diameter) which changed color at randomized intervals between 1 and 5 seconds. Subjects were asked to fixate on the dot and indicate each time the dot changed color using an MRI compatible button box. The stimuli were presented through either wedge, ring, or bar apertures and consisted of achromatic contrast patterns (spatially pink noise) overlaid with randomly positioned and scaled visual object stimuli from Kriegeskorte et al. (2008). The full stimuli patterns can be found at http://cvnlab.net/analyzePRF/.

In the first set of experiments the stimulus consisted of a combination of rotating wedge, contracting ring and sweeping bar stimuli. The stimuli can be broken down into the following temporal structure:

- Wedge stimuli covering a 90° angle swept 8 full counter-clockwise rotations. The wedge took 32 seconds to complete one full 360° rotation.
- Bars with a width equal to 12% of the stimulus diameter were swept across 8 unique (cardinal and oblique) directions. Each bar sweep took 32-seconds to complete (28-second sweep followed by a 4-second rest).
- Contracting ring stimuli were swept across the visual field over the course of 32-seconds (28-second sweep followed by a 4-second rest). This was repeated 5 times. The widths of the rings were scaled linearly with eccentricity to compensate for cortical magnification in visual cortex (Cowey and Rolls, 1974; Rovamo and Virsu, 1979).
- 16 second rest periods with mean luminance (zero contrast) images were placed at the start, between stimuli types and at the end of the stimulus (taking into account the additional 4 second rest periods at the end of bar and ring sweeps).

The stimulus took a total of 12 minutes 20 seconds to complete (16 + 8^*^32 + 16 + 4^*^32 + 12 + 4^*^32 + 12 + 5^*^32 + 12 s). We shall refer to simulation and in *in-vivo* data which used these stimuli as Dataset 1.

In the second set of experiments we investigated the robustness of model fitting for stimuli of different lengths. Each of the 4 runs consisted of the same 5 minute sweeping bar stimuli. The temporal structure was as follows:

- 5 minute sweeping bar stimuli: Bar width was set to 12% of the stimulus diameter. Temporal
structure was organized as 16 seconds of mean luminance rest followed by 4 bar sweeps of 32 seconds duration. 12 seconds of mean luminance rest followed by 4 additional bar sweeps of 32 seconds duration. Then a final 16 seconds of mean luminance rest.

We refer to experiments using these stimuli as Dataset 2.

### Simulations

Synthetic data was generated using the CSS-pRF model of Kay et al. (2013). A total of 512 different receptive fields were tested using combinations of parameters typical of pRF properties within human visual field maps. To avoid any potential bias between the estimation methods we chose a set of simulation ground-truth values distinct from those explicitly modeled in the CSS_CO-pRF_ dictionary (*n*=1 being the exception). The following parameters were used: polar-angle *θ* = {0,45,90,135,180,225,270,315}, eccentricity *r* = {2.5,13.3,24.2,35}, pRF size *σ* = {2.5,10,17.5,25}, exponent *n* = {0.15,0.43,0.72,1} and gain *g*=5. The parameters *r* and *σ* are defined as a percentage of total stimulus coverage (e.g. across the stimulus diameter). The signal was then contaminated with Gaussian noise to provide a specified contrast-to-noise ratio (CNR) between 3 and 10. Our calculation of CNR follows definition 2 of Welvaert and Rosseel (2013):

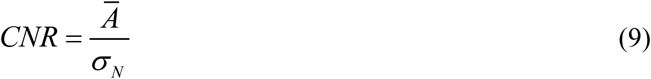

where *A* is the mean across all peak amplitudes in a simulated response and *σ_N_* is the standard deviation of the noise. For each receptive field we modeled 20 unique noise realizations resulting in a total of 10240 experiments for each *CNR.* All simulations assumed a canonical hemodynamic response function.

The simulations were divided into two sets of experiments. The first experiments tested the simulated receptive field responses using the 12 min 20 second combined wedge, bar and ring stimuli of Dataset 1. The second set of experiments investigated parameter estimation when using the 5 minute sweeping bar stimuli of Dataset 2. These 5 minute stimuli runs were repeated either 1 or 4 times resulting in datasets with total stimuli durations of 5 or 20 minutes.

### In-vivo datasets

All MRI data was acquired with a 3T Siemens Prisma MR system using a 64-channel head coil. Two male subjects with no history of eye disease (aged 28 and 30 yrs) underwent all of the fMRI acquisition protocols. Foam padding and audio headphones minimized head motion.

The functional MRI data were acquired using a 2D-EPI sequence (TR/TE = 1518/30 ms, 23 slices, 3x3x2.5 mm^3^ resolution, 20% distance factor, FOV = 192x192 mm^2^, 12% phase oversampling). The slices were oriented parallel to the calcarine sulcus, covering the majority of occipital cortex. A whole brain 2D-EPI sequence with 49 slices and 5 volumes was acquired to aid with structural registration. All other parameters of the whole brain sequence were the same as in the fMRI acquisition. A T1-weighted anatomical image (TR/TE = 2000/2.93 ms, 1x1x1 mm^3^ resolution) was acquired and used to reconstruct the cortical surfaces. In order to estimate local field distortions we acquired a B0 field map (TR/TE = 1020/10 ms, 3x3x3 mm^3^ resolution).

Functional scans for Dataset 1 used the combined rotating wedge, contracting ring and sweeping bar stimuli and were acquired over 488 volumes. A total of 3 fMRI scans were performed in the session and later used to investigate scan-rescan reproducibility of the estimated pRF parameters. In a separate session functional scans for Dataset 2 were acquired using the 5 minute sweeping bar stimuli and 203 fMRI volumes. The 5 minute fMRI acquisitions were repeated over 4 runs and later used to assess parameter estimation for 5 minutes versus 20 minutes of data.

### Image preprocessing

The first 5 volumes of each fMRI run were discarded to allow magnetization to reach a steady state. Preprocessing of the fMRI data was performed within SPM12 using the following procedure. Slice-time correction was applied to adjust for differences in slice acquisition times. The fMRI data was aligned to correct for movement artifacts and co-registered with the whole brain reference data. The fMRI and reference data were spatially unwarped (Andersson et al., 2001; Jezzard and Balaban, 1995) to improve coregistration between the EPI and structural data (Hutton et al., 2002). The reference data was then coregistered with the T1-weighted structural image and the same rigid-body transformation applied to the fMRI data. Binary stimuli masks were defined at time points which coincided with the slice-time corrected fMRI data and down sampled to a resolution of 100x100 pixels to improve efficiency of the nonlinear optimization routines.

The T1-weighted anatomical images were processed with FreeSurfer (Dale et al., 1999) for white and grey matter segmentation and cortical surface reconstruction. FreeSurfer’s Desikan-Killiany atlas (Desikan et al., 2006) was used to select cortical regions within the occipital lobe and to reduce to the amount of cortex over which we fit the pRF models. The preprocessed fMRI data was then sampled onto the FreeSurfer surface representations at a depth 20% from the white matter interface. All CSS-pRF model estimations were performed on the surface projected fMRI responses. Visual field delineation was performed manually using the polar angle maps projected onto inflated cortical surfaces (Wandell et al., 2007).

### pRF model estimation

The parameters of the CSS-pRF model (Dumoulin and Wandell, 2008; Kay et al., 2013a) were estimated using both the CSS_CO-pRF_ formulation described within this work and the CSS_Orig_ nonlinear fitting procedure typical of current approaches within the literature.

To solve Eq. (6) efficiently for CSS_CO-pRF_ we used an implementation of the LASSO algorithm (Tibshirani, 1996) within the *SPArse Modeling Software* (SPAMS; http://spams-devel.gforge.inria.fr/) Matlab toolbox. Prior to solving Eq. (6), polynomial regressors were used to detrend the in-vivo fMRI data and the CSS_CO-pRF_ dictionary responses. A separate set of polynomial regressors were used for each run with a maximum degree of 6 for Dataset 1 and maximum degree of 3 for each run of Dataset 2. Detrended dictionary responses were then normalized using the *l*_2_-norm. These procedures removed low frequency artifacts and ensured an optimal model fit.

The CSS_Orig_ procedure is based on the methods within Kay et al. (2013), details of which and the code are available at http://cvnlab.net/analyzePRF/. The fitting procedure uses a Levenberg-Marquardt algorithm for optimization, minimizing squared error between the CSS-pRF model and the data. An initial seed location is chosen from a simple grid search similar to the method proposed by Dumoulin and Wandell (2008) as well as two additional seeds of different receptive field sizes centered with respect to the stimulus. A two-stage optimization strategy is used for each seed whereby all parameters excluding the exponent parameter are first optimized (holding the exponent parameter fixed) and then all parameters are optimized (including the exponent parameter with an initial seed value *n* = 0.5). This strategy was motivated by the need to reduce local minima issues when using a non-linear optimization.

All model estimations were performed on a single core of a DALCO high-performance cluster without multi-threading or parallel computing.

## Results

### Simulation results: Dataset 1

Fig. 1 evaluates CSS-pRF parameter estimation for CSS_CO-pRF_ and CSS_Orig_ as a function of CNR over the 512 simulated receptive field responses in Dataset 1. The performance of parameter estimation for polar-angle *θ*, eccentricity *r*, pRF size *σ*, exponent *n* and gain *g* were assessed via the absolute error and correlation with known ground-truth values at each CNR value. The results validate the CSS_CO-pRF_ framework as providing accurate and precise parameter estimates. Across all tested CNR values and model parameters the CSS_CO-pRF_ estimates had significantly less absolute error as compared with the equivalent CSS_Orig_ estimates (two-sample paired sign test, p < 0.001 FDR corrected). Furthermore the CSS_CO-pRF_ estimates for *r*, *σ*, *n* and *g* across all tested CNR levels had significantly less absolute error than the CSS_Orig_ estimates in the absence of noise (CNR = ∞; two-sample paired sign test, p < 0.001 FDR corrected).

**Fig. 1.**
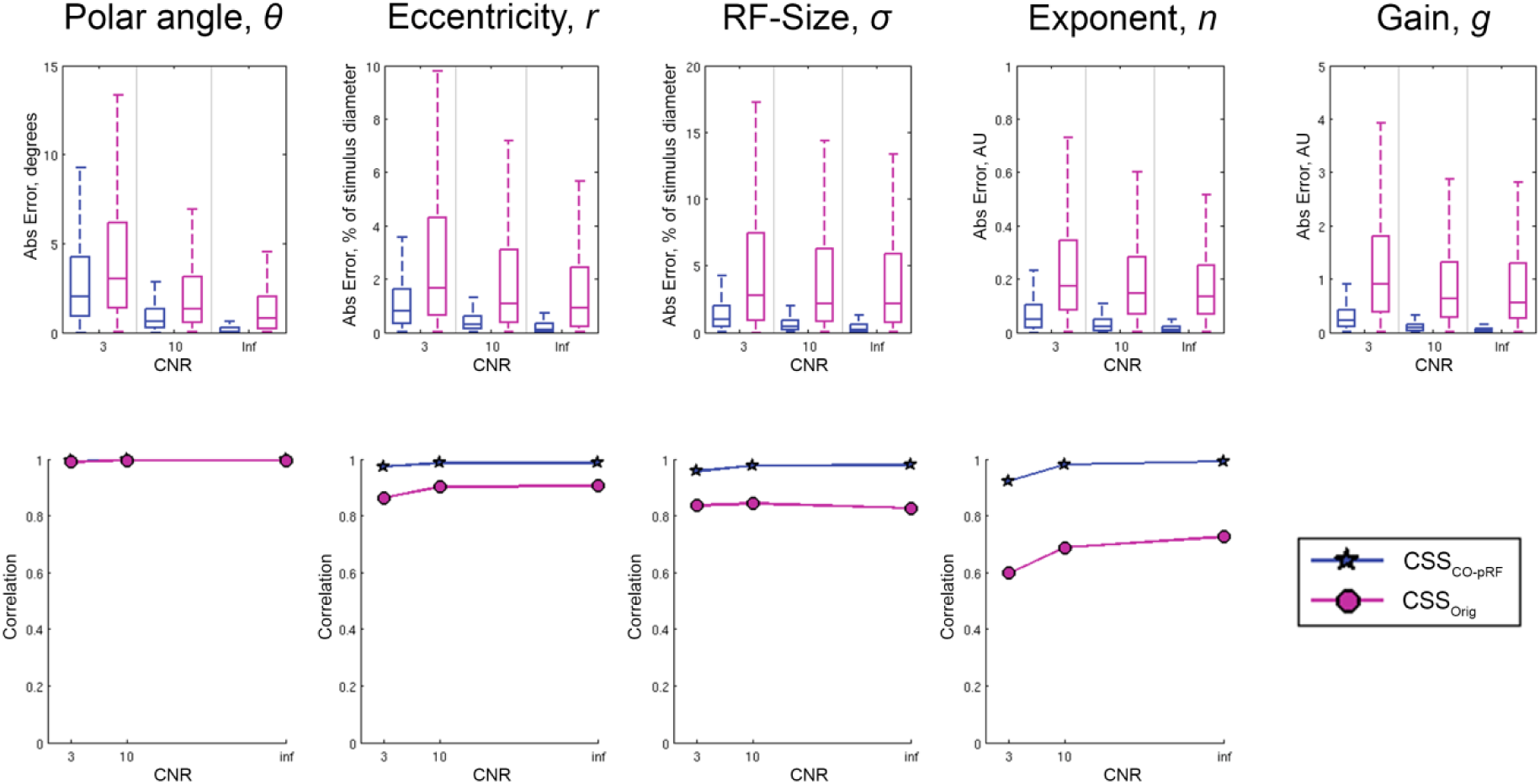
CSS-pRF parameter estimation performance on synthetic data as a function of CNR. The model parameter estimates of the CSS_CO-pRF_ algorithm (blue) are compared with those from the CSS_Orig_ algorithm (purple) by means of the absolute error (top) and the correlation of the estimated parameters with respect to the known ground-truth (bottom). The whiskers of the absolute error box plots extend to the outermost data points within 1.5 times the interquartile range.

The correlations with ground-truth values suggest that CSS_CO-pRF_ results in parameter estimates with reduced bias as compared to CSS_Orig_. Both approaches produce robust ground-truth correlations for estimates of **θ** however CSS_CO-pRF_ yields a stronger correlation than CSS_Orig_ for *r*, *σ* and *n* across the full range of investigated CNR values. This is demonstrative of the inherent bias of nonlinear optimization routines due to the choice of initial seed parameters and local minima issues. We do not present estimated versus ground-truth correlations for *g* as all simulations had the same ground-truth value. The first column of Table 1 details the computational time required to fit both algorithms to the simulated responses of Dataset 1. The CSS_CO-pRF_ fit considerably reduces the computational time by a factor of 47 as compared to the CSS_Orig_ estimation.

**Table 1.** Computational times for the CSS_CO-pRF_ and CSS_Orig_ algorithms on simulation and in-vivo data. The CSS_CO-pRF_ algorithm accelerates model fitting by a factor of 50 (approx.).

|  | Simulations: Dataset 1 |  | In-vivo data |  |
| --- | --- | --- | --- | --- |
|  | Total time | Time per voxel | Total time | Time per voxel |
| $\text{CSS}_{\text{Orig}}$ | 03 days 02 hrs 45 min | 6.57 s | 20 days 23 hrs 16 min | 6.44 s |
| $\text{CSS}_{\text{CO-pRF}}$ | 01 hrs 35 min 34 s | 0.14 s | 10 hrs 09 min 46 s | 0.13 s |

The data in Fig. 2 expands upon the simulation results for the cases of zero noise and CNR = 3. Estimated parameters are presented as box and whisker plots for CSS_CO-pRF_ (blue) and CSS_Orig_ (purple) algorithms across all parameter ground-truth values. These results confirm the high precision and accuracy of parameter estimation for the CSS_CO-pRF_ approach across individual model parameter values and noise levels.

**Fig. 2.**
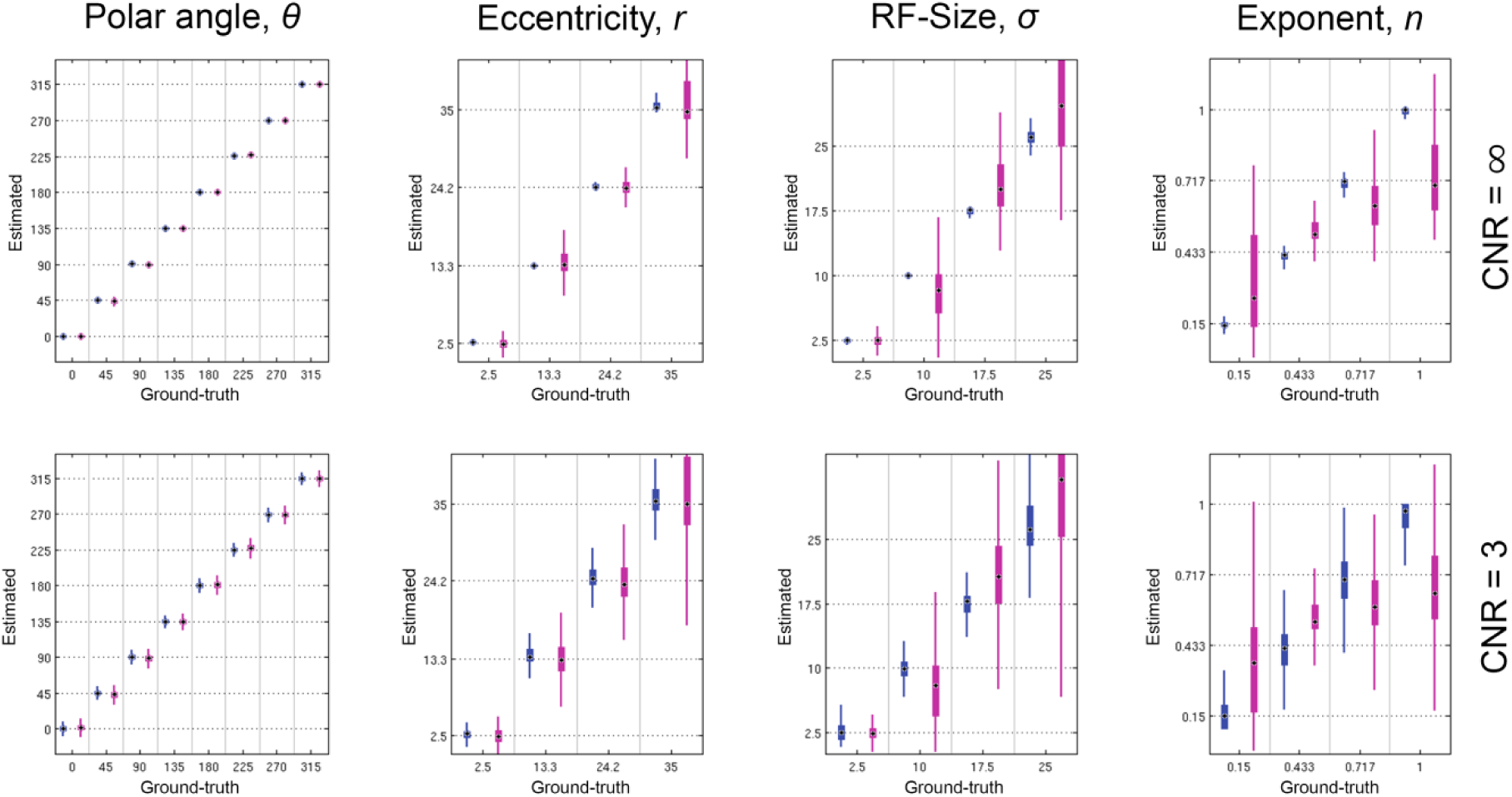
Detailed evaluation of CSS-pRF parameter estimation on synthetic data. The model parameter estimates from the CSS_CO-pRF_ algorithm (blue) are compared with those from the CSS_Orig_ algorithm (purple) versus ground-truth values for noise free (top) and CNR = 3 (bottom) data. Ground truth values are indicated with dashed horizontal lines. The whiskers of the box plots extend to the outermost data points within 1.5 times the interquartile range. Median values are indicated with a black target.

Both methods perform more or less equivalently at estimating *θ* while CSS_Orig_ has reduced precision when estimating *r* and reduced accuracy and precision when estimating *σ* or *n*. For both the noise-free and CNR = 3 data CSS_Orig_ tends to overestimate *σ* with increasing eccentricity and displays a reduced dynamic range for *n* centered on 0.5. This is a further demonstration of the seeding bias and local minima issues for the CSS_Orig_ nonlinear optimization routine as parameter estimates lie close to the initial seed value of *n* = 0.5. CSS_CO-pRF_ has reduced precision in the presence of noise but the parameter estimates remain accurate and unbiased. Furthermore a comparison of the noisy CSS_CO-pRF_ estimates (bottom row blue) with the noise free CSS_Orig_ estimates (top row purple) suggests that the CSS_CO-pRF_ algorithm achieves comparable parameter estimates with noisy data as compared to the CSS_Orig_ algorithm with noise free data.

### In-vivo results: Dataset 1

Fig. 3 compares the estimated in-vivo parameter maps of Dataset 1 for the CSS_CO-pRF_ and CSS_Orig_ algorithms on a representative inflated left hemisphere. The standard deviation of parameter estimates across the three repeated acquisitions is also provided. The maps of polar angle, *θ*, appear almost identical between the two approaches, both in terms of the identification of visual field map boundaries and the reproducibility of estimates across runs. The CSS_Orig_ algorithm tends to produce elevated eccentricity, *r*, and rf-size, *σ*, estimates in regions responding to peripheral areas of the visual field (<9° of visual angle). These regions also exhibit increased CSS_Orig_ parameter variance across the three runs. Both methods provide largely comparable estimates for *r* and *σ* in areas responding to more central portions of the visual field (<9° of visual angle). One difference of note between the two methods is in the estimation of the exponent, *n*, parameter. CSS_CO-pRF_ suggests a greater level of variation across visual cortex compared to the CSS_Orig_ estimates which are centered on 0.5 with reduced dynamic range. The observed differences in the estimation of *n* likely result from seeding and local minima issues of the nonlinear algorithm and reflect the simulation results where the ground-truth is known. Across all parameter maps CSS_CO-pRF_ produces smoother appearing maps than those produced by CSS_Orig_.

**Fig. 3.**
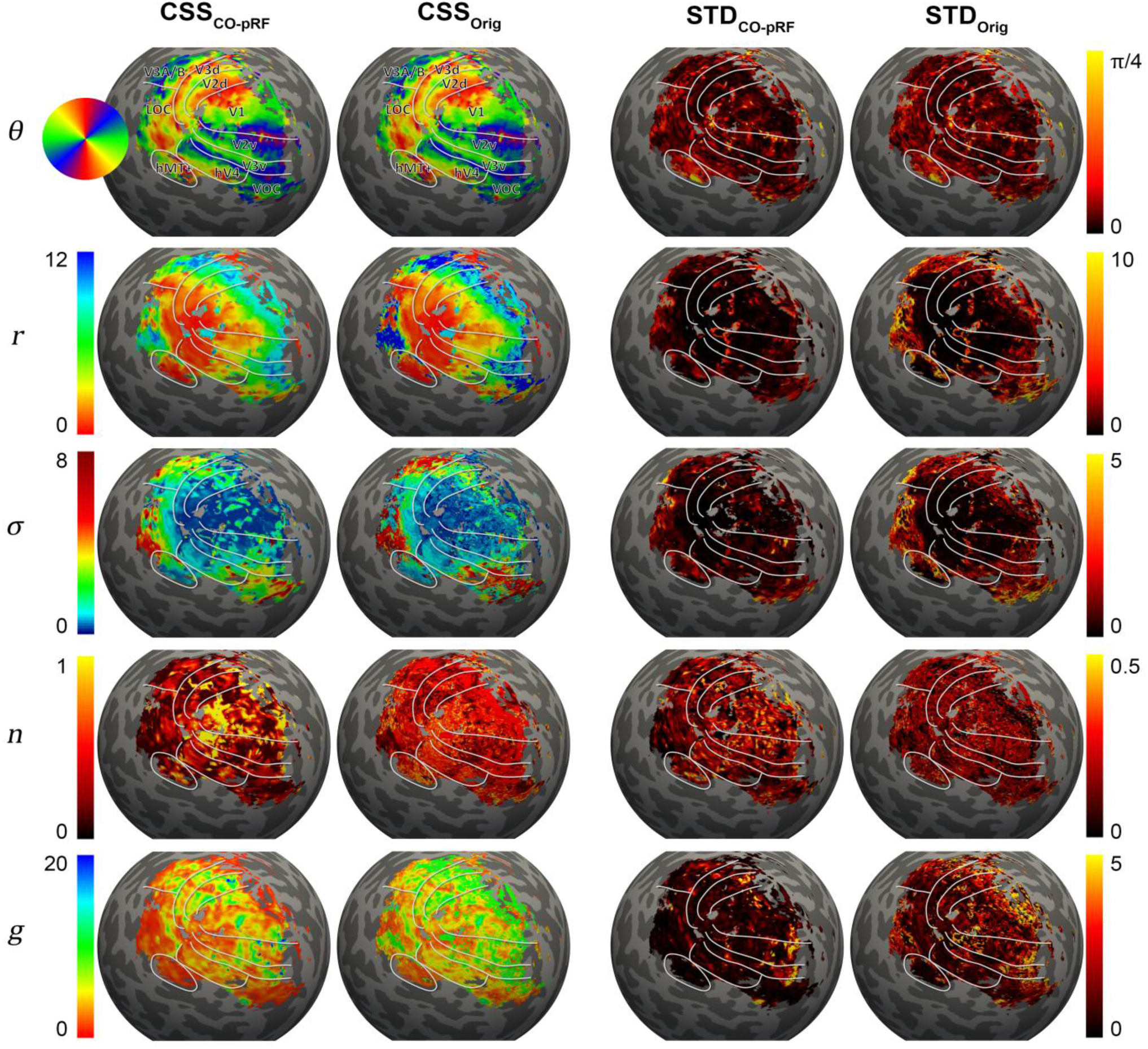
Model parameter maps for in-vivo dataset 1. The CSS-pRF parameters *θ*, *r*, *σ*, *n* and *g* estimated using the CSS_CO-pRF_ and CSS_Orig_ algorithms are reported on the inflated left hemisphere of a representative subject. The last two columns show the standard deviation of parameter estimates across the three repeated acquisitions.

An assessment of explained variances between the two methods can be seen in Fig. 4. In Fig. 4A the differences in explained variance
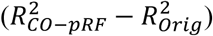
are displayed on an exemplary inflated left hemisphere. We only plot vertices where both
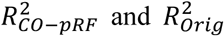
are greater than 0.05. The CSS_CO-PRF_ method systematically outperforms CSS_Orig_ by explaining more variance across responsive visual field areas. This is further revealed in the scatter plot of Fig. 4B which compares the coefficients of determination (*R*^2^) for the final CSS-pRF parameter estimates. The *R*^2^ estimates are highly correlated although the CSS_CO-pRF_ algorithm gets closer to the global minima in 93% of the fit responses, on average explaining 2% more of the variance. This systematic difference in explained variance provides strong evidence that CSS_CO-pRF_ results in parameter estimates closer to the global optimum solution.

**Fig. 4.**
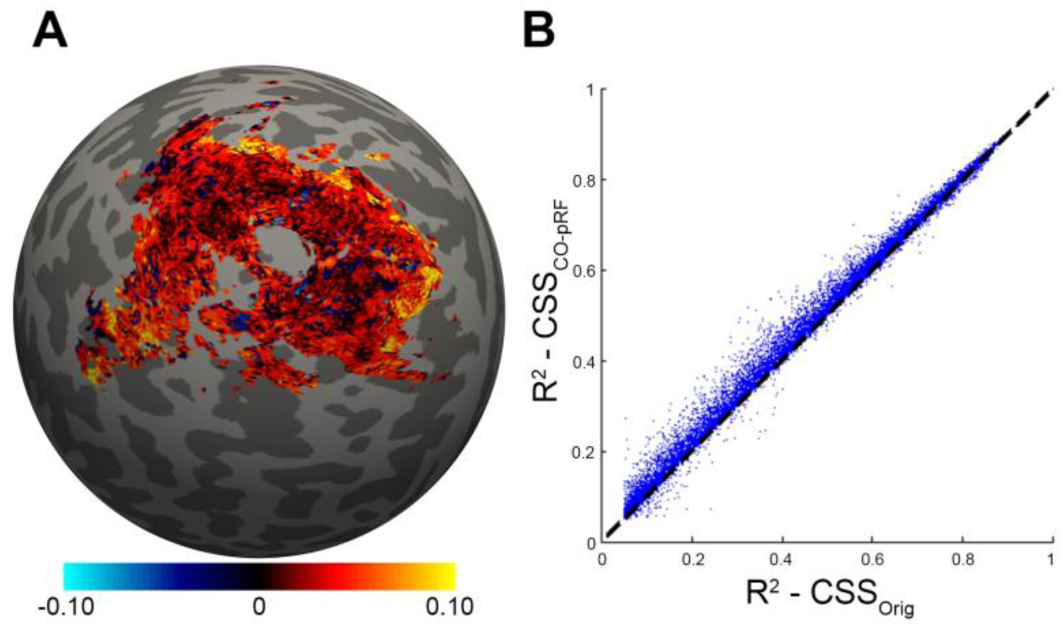
Coefficient of determination (*R*^2^) comparisons. The difference in explained variance of the CSS_CO-pRF_ and CSS_Orig_ formulations is displayed on an inflated left hemisphere (A). A scatter plot of *R^2^-CSS_CO-pRF_* versus *R*^2^-*CSS_Orig_* for estimated fMRI responses across the surface (B). The CSS_CO-pRF_ approach explains more of the measurement variance across 93% of the fit responses despite estimating the parameters in a fraction of the computational time. Only vertices with *R*^2^-*CSS_CO-pRF_* and *R*^2^-*CSS_Orig_* greater than 0.05 are displayed.

Table 1 shows the computational time required to fit both algorithms to the in-vivo acquisitions. The second column corresponds to the data presented in Fig. 3 and Fig. 4. The CSS_Orig_ algorithm took approximately 21 days to fit the model to all of the selected fMRI responses in occipital cortices, corresponding to an average fit time of 6.44 s/response. Conversely the CO-pRF approach drastically reduced the computational burden, estimating the same set of fMRI responses in approximately 10 hrs, just 0.13 s/response. The increased computational efficiency afforded by the convex reformulation resulted in an acceleration factor of 50 for the in-vivo processing of Dataset 1.

### Simulation results: Dataset 2

CSS-pRF parameter estimates for the CSS_CO-pRF_ and CSS_Orig_ algorithms are presented as a function of stimulus duration and CNR in Fig. 5. As in Fig. 1 the parameter estimation performance for polar-angle *θ*, eccentricity *r*, pRF size *σ*, exponent *n* and gain *g* were assessed via the absolute error and correlation with known ground-truth values. The results suggest that CSS_CO-pRF_ provides robust parameter estimates in the case of reduced data. Across all tested CNR levels and stimulus durations the key pRF parameters *θ*, *r* and *σ* CSS_CO-pRF_ have significantly less absolute error than the equivalent CSS_Orig_ estimates (two-sample paired sign test, p < 0.001 FDR corrected). Furthermore the 5 minute CSS_Orig_ estimates for *r* have significantly less absolute error than the 20 minute CSS_Orig_ estimates across all SNR values (two-sample paired sign test, p < 0.001 FDR corrected). The CSS_Orig_ parameter estimates also tend to have a reduced correlation to known ground-truth regardless of stimulus duration or noise level. Both algorithms perform comparatively poorly when estimating *n* or *g*; the reduced estimation precision in contrast to the results presented in Fig. 1 suggest that the chosen stimulus type (sweeping bars vs. combined wedges, rings and bars) significantly influences the estimation of *n* and *g*.

**Fig. 5.**
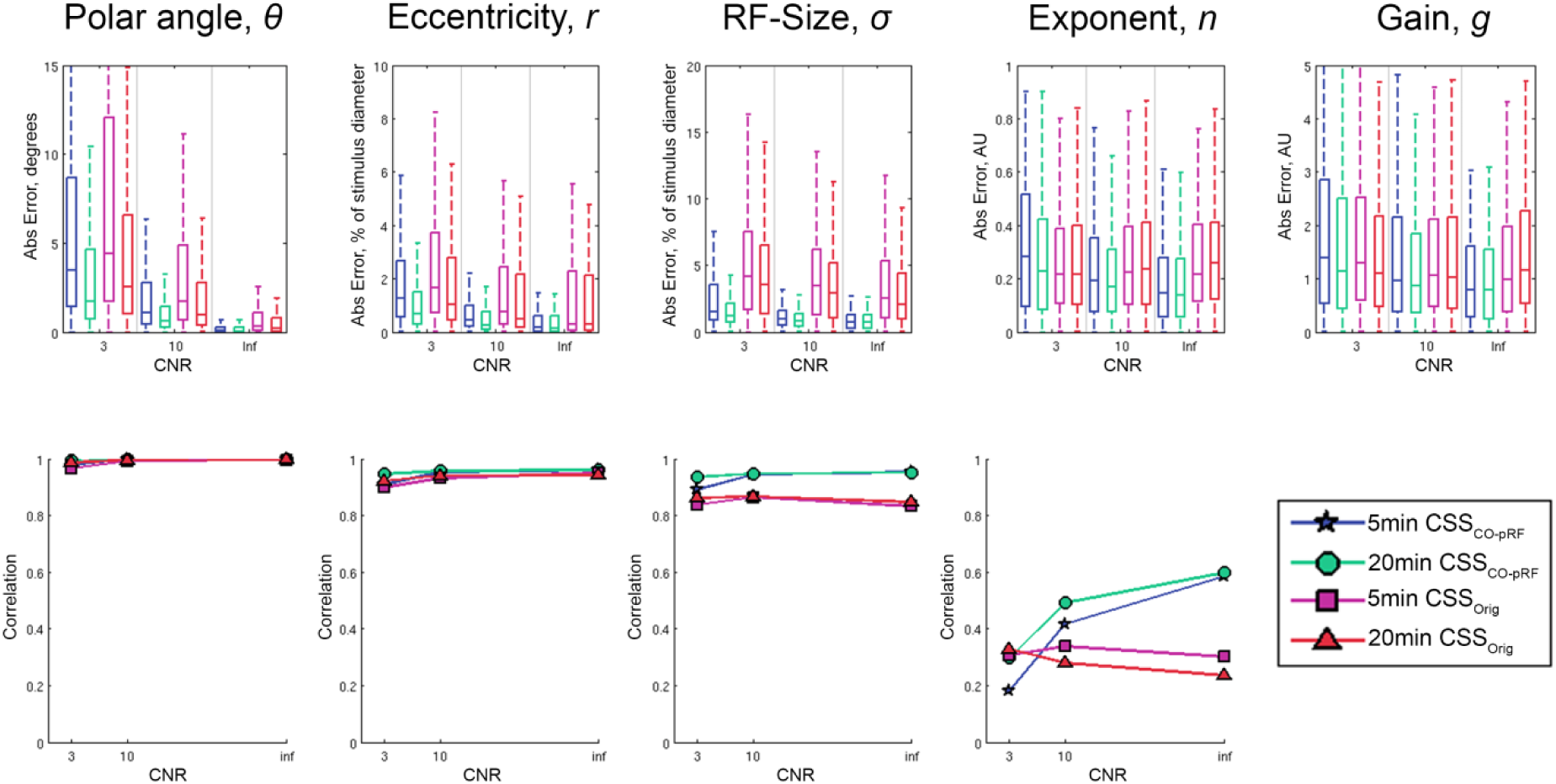
CSS-pRF parameter estimation performance on synthetic data as a function of stimulus duration and CNR. The model parameter estimates of the CSS_CO_-_pRF_ algorithm with 5 minute (blue) and 20 minute (green) stimulus duration are compared with those from the CSS_Orig_ algorithm with 5 minute (purple) and 20 minute (red) stimulus duration by means of the absolute error (top) and the correlation of the estimated parameters with respect to the known ground-truth (bottom). The whiskers of the absolute error box plots extend to the outermost data points within 1.5 times the interquartile range.

Fig. 6 expands upon the stimulus duration results for the cases of zero noise and CNR = 3. Both methods are capable of providing accurate and precise estimates for *θ* regardless of stimulus duration or noise level. For both methods the parameters *r* and *σ* have reduced precision with the addition of noise and/or reduced stimulus duration however CSS_CO-pRF_ remains accurate while there is some bias for the CSS_Orig_ estimates of *σ*. Moreover the 5 minute CSS_CO-pRF_ estimates (blue) for *r* and *σ* compare favorably to the 20 minute CSS_Orig_ estimates (red).

**Fig. 6.**
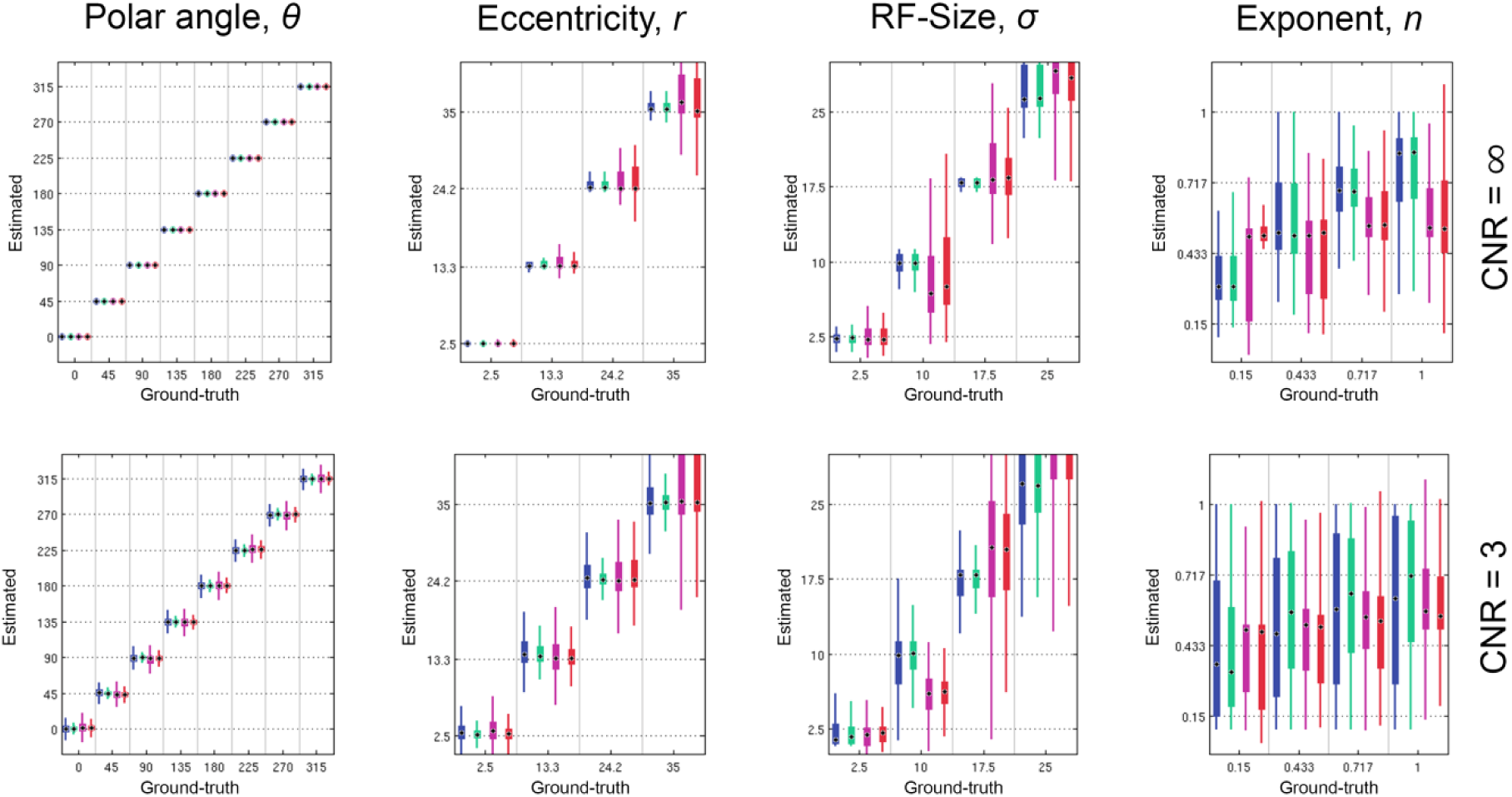
Detailed evaluation of CSS-pRF parameter estimation on synthetic data with varying stimulus duration. The model parameter estimates from the CSS_CO-PRF_ algorithm with 5 minute (blue) and 20 minute (green) stimulus duration are compared with those from the CSS_Orig_ algorithm with 5 minute (purple) and 20 minute (red) stimulus duration versus ground-truth values for noise free (top) and CNR = 3 (bottom) data. Ground truth values are indicated with dashed horizontal lines. The whiskers of the box plots extend to the outermost data points within 1.5 times the interquartile. Median values are indicated with a black target.

### In-vivo results: Dataset 2

A comparison of *σ* maps estimated with either 5-minutes or 20-minutes of fMRI data is shown in Fig. 7. The CSS_CO-pRF_ algorithm produces smooth and reproducible maps even for short acquisition durations. In contrast CSS_Orig_ estimates are noisier as can be seen in the ratio of *σ* estimates between the two acquisition durations *σ*_20min_/*σ*_5min_. The ground truth *σ* values are unobtainable for in-vivo fMRI measurements. However, for the 20 minute data acquisition the CSS_CO-pRF_ fit explained more of the measurement variance than CSS_Orig_ across 92.5% of vertices. Furthermore the estimated model parameters fit using CSS_CO-PRF_ using 5 minutes of data explained more of the measurement variance than the CSS_Orig_ estimates fit using 20 minutes of data across 56.2% of the vertices. This suggests that CSS_CO-pRF_ is robust under reduced data conditions while also being more successful than CSS_Orig_ at achieving a set of parameter estimates close to the global optimum at equivalent and even reduced data conditions.

**Fig. 7.**
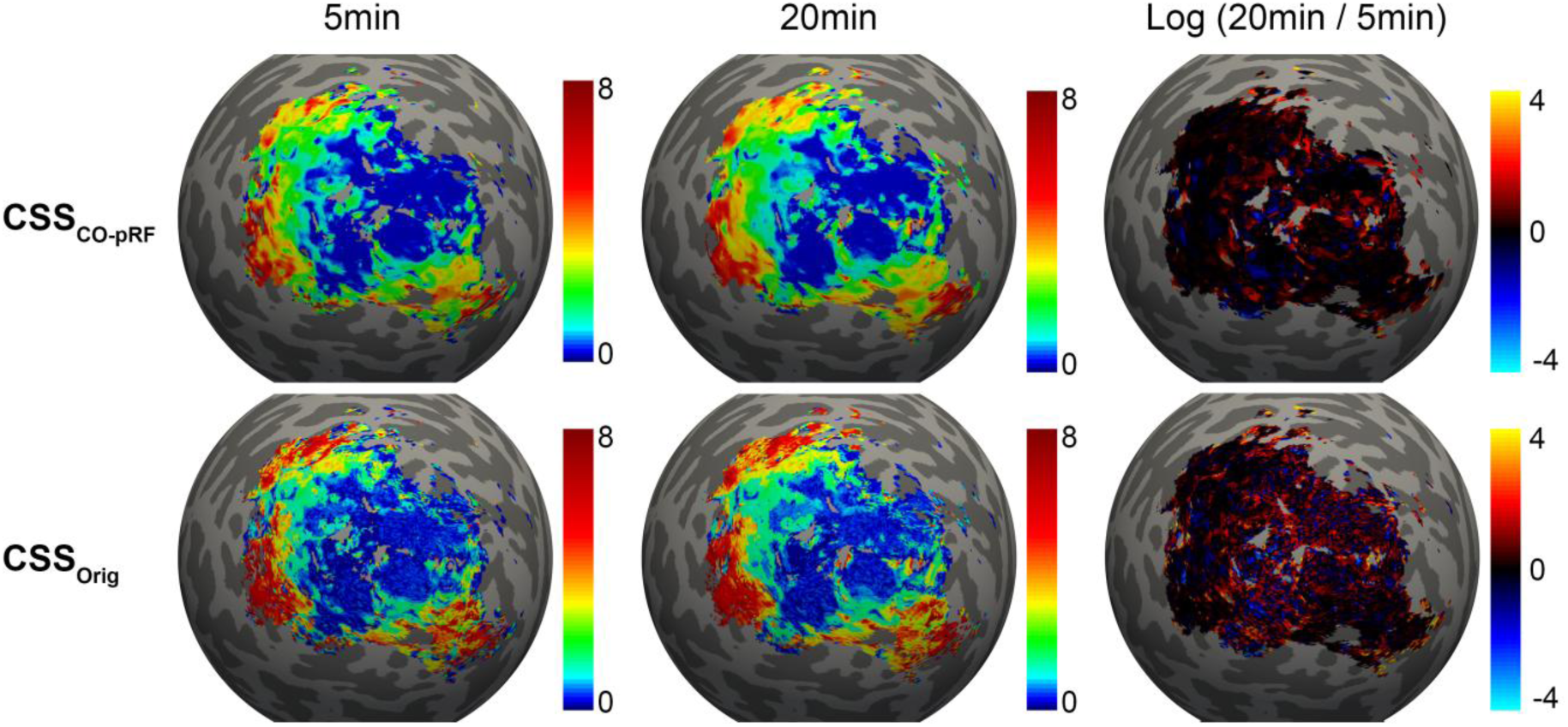
Estimation of population receptive field sizes (*σ*) for short (5-minute) and longer (20-minute) fMRI acquisition durations using the CSS_CO-pRF_ and CSS_Orig_ algorithims. The CSS_CO-pRF_ estimates produce smooth and consistent maps for both 5 minutes and 20 minutes of acquired data.

## Discussion

### Advantages and limitations

The principle aim and benefit of the CO-pRF framework is to accelerate model parameter estimations by an order of magnitude or greater while achieving model parameter estimates close to the global optimal solution with little or no bias. A number of studies have begun to apply pRF methods to the investigation of visual function in clinical populations (Clavagnier et al., 2015; Papanikolaou et al., 2014; Schwarzkopf et al., 2014); the introduction of faster CO-pRF algorithms (up to a factor of 50 or more) may result in the use of pRF methods on a larger scale, without the need for powerful supercomputing clusters. Furthermore our results suggest that CSS_CO-pRF_ is robust even in the case of reduced scan time, with 5 minutes of fMRI data providing CO-pRF parameter estimates broadly comparable to those produced using current methods with 20 minutes of data. The possibility of reducing required scan durations may further enhance the applicability of pRF methods to situations in which typical fMRI scan times are not feasible.

In addition to greatly accelerating the model fit our simulation and in-vivo results suggest that CSS_CO-pRF_ can achieve accurate and robust parameter estimates with reduced bias and variance as compared to CSS_Orig_. The CSS_Orig_ algorithm uses a two-stage fitting procedure to estimate model parameters whereby a dense grid search is used to select putative seed values, followed by a Levenberg-Marquardt non-linear optimization routine on these initial seed parameters and two additional seeds centered with respect to the stimulus. CSS_Orig_ then selects which of the three non-linear optimizations reduces residual error the most and saves the corresponding parameter estimates. In comparison the convex reformulation of CSS_CO-pRF_ linearizes the optimization procedure and provides a sparse estimate of weighted dictionary elements whose averaged underlying parameters are used to predict the data. One might conceive of this procedure as an interpolation of dictionary elements clustered around the global minima solution. Non-linear optimization procedures are particularly susceptible to getting trapped in local minima of the objective function, especially if the model parameters are correlated as has been suggested to be the case for visual pRF models (Zeidman et al., 2016). Bias due to local minima issues for the CSS_Orig_ procedure is demonstrated by the reduced ground truth correlations of Fig. 1 for simulated data and a reduction to explained variance for in-vivo data as shown in Fig. 4. In contrast, the CO-pRF formulations are convex and thus will guarantee convergence to the global minima solution supported by the dictionary.

One area of consideration for CO-pRF formulations is the way in which the dictionary is constructed. It is important to select underlying parameters which cover the full range of biologically meaningful measurements. However aside from this consideration a number of choices can be made to tradeoff the estimation precision of the different parameters against the computational time required for the model fit. For example, one might like to improve the estimation of receptive field sizes, *σ*, in a certain study, perhaps because of a hypothesized difference in *σ* between two groups of individuals. This can be easily achieved by simply increasing the number of values for *σ* used to create the CO-pRF dictionary at the expense of some computational time as the full dictionary size correspondingly increases. Despite the importance and potential benefits of exploiting dictionary construction methods a detailed investigation of these goes beyond the purpose of the present work, this being to provide evidence that robust, unbiased parameter estimates can be achieved within a fraction of the computational time required by current methods.

### Applications and future directions

The results of this paper provide a proof of principle for the application of the CO-pRF framework to problems of sensory receptive field mapping. Across both synthetic and real data experiments CO-pRF estimates were accelerated by greater than an order of magnitude whilst being robust and in good agreement with original methods. This therefore achieves the goal of the present work.

In this work we focused our attention on the CSS-pRF model of visual cortex responses (Kay et al., 2013a). This method assumes a circularly symmetric Gaussian receptive field shape however more complicated shapes have been suggested within the literature (Greene et al., 2014; Kay et al., 2008; Lee et al., 2013; Sprague and Serences, 2013; Zuiderbaan et al., 2012). Other work has extended the pRF model to include second-order contrast information, taking into account stimulus patterns as well as stimulus locations (Kay et al., 2013b). Furthermore, although our example application assumed a canonical HRF form one might like to apply a parameterized model of neuronal activation to blood flow coupling (e.g. Buxton and Frank, 1997; Friston et al., 2000). In each case the flexibility of the CO-pRF framework would allow these extensions to be incorporated into the linear systems of Eq. (6). In fact multiple model variants could be combined in full and modeled within the same partitioned dictionary. A model selection procedure, such as the Bayesian information criterion, could then be used to select the model type which is best supported by the data. Nonetheless, for the sake of clarity, within this work we chose to focus on the introduction of a generalized framework followed by an example application to a single pRF model. We will investigate more complicated pRF model extensions, their combination and model selection procedures within future work.

CO-pRF estimates are fast and robust; however there may be circumstances in which another fitting procedure is advantageous. A recent work proposed a Bayesian estimation of pRF model parameters (Zeidman et al., 2016) and in such a case the CO-pRF algorithm could be run quickly to provide the initial seed estimates required as input to the procedure. A Bayesian estimation approach has the advantage of providing estimates of measurement uncertainty; however it is unlikely to be applicable in all cases as the approx. 100 second fit time per voxel would make a Bayesian estimation of large datasets extremely challenging even if processed on high performance computing clusters. Another method which could be used following the CO-pRF estimation is that of Markov chain Monte Carlo (MCMC). Typically these approaches are computationally slow, however work by our group to develop a high performance implementation using a Graphics Processing Unit (GPU) (Adaszewski et al., 2016) has proved promising and in the future we intend to integrate the GPU optimized MCMC procedure within the CO-pRF toolbox.

Our example application of the generalized CO-pRF framework to a visual pRF model was motivated by the fact that these models are well established within the field of visual neuroscience (Dumoulin and Wandell, 2008; Wandell and Winawer, 2015) and follow on from decades of research into the retinotopic organization of human visual cortex (DeYoe et al., 1996; Engel et al., 1994; Sereno et al., 1995). Recent developments within the field of human auditory neuroscience offer the potential to quantitatively map auditory processes non-invasively using the pRF methodology (Thomas et al., 2015) and would be well suited to being solved within the CO-pRF framework. Beyond the visual and auditory system one might even conceive of analogous pRF mapping methodologies being developed for somatosensory, olfactory or gustatory cortices provided appropriate stimuli, computational models and recording techniques are available. Indeed pRF methodologies need not be restricted to fMRI techniques and merely require a model of modality specific responses to a neural event. For example pRF techniques might well be extended to measurements made using EEG/MEG by applying a neural mass model (David et al., 2005; David and Friston, 2003) of neuronal signal coupling in place of the hemodynamic response function used in fMRI.

## Conclusions

This study aimed to improve upon the long and often impractical fitting times required by existing population receptive field techniques. By reformulating the pRF equations in a fashion solvable by convex estimation procedures the CO-pRF framework offers the potential to vastly improve model fitting times while achieving superior model parameter estimates, closer to the global minima solution. The CO-pRF framework is flexible and capable of estimating receptive fields of arbitrary shape, sensory modality and measurement techniques. We chose the CSS-pRF fMRI technique as an example methodology for comparison. Across a wide range of measures CO-pRF fitting achieved model parameter estimates closer to the global optimum while requiring only 1/50^th^ of the computational time. By making available such ultrafast pRF fitting algorithms it becomes increasingly practical to apply pRF methods to large cohorts of patients and to the longitudinal assessment of pRF measures over time. With this in mind we provide a Matlab based toolbox for CO-pRF model fitting (https://github.com/davesl/COpRF) and hope that these tools prove useful for the sensory neuroscience community.

## Acknowledgements

The research leading to these results has received funding from the European Union Seventh Framework Programme (FP7/2007-2013) under grant agreement No 604102 and the European Union’s Horizon 2020 research and innovation programme under grant agreement No 720270 (HBP SGA1). BD is supported by the Swiss National Science Foundation (NCCR Synapsy, project grant Nr 32003B_159780 and SPUM 33CM30_140332/1), Foundation Parkinson Switzerland and Foundation Synapsis. This work was carried out on the MRI platform of the Département des Neurosciences Cliniques - Centre Hospitalier Universitaire Vaudois, which is generously supported by the Roger de Spoelberch and Partridge Foundations. We would like to thanks the LREN group for helping in the acquisition of the MRI data used in this paper.

